## Supplementary figures and images for "Positional SHAP (PoSHAP) for Interpretation of Machine Learning Models Trained from Biological Sequences"

### SuppFigure_3.png

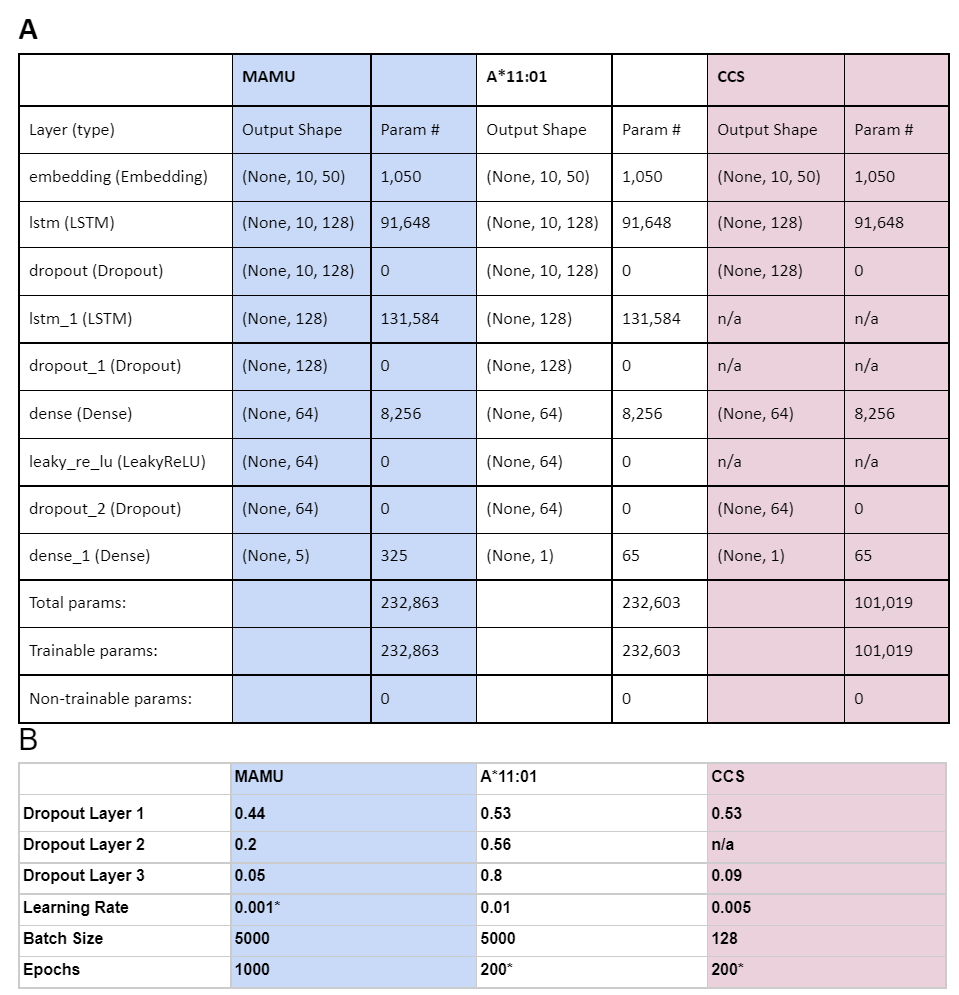
